# Comparative analysis of sfRNA in the genome of pre-epidemic and epidemic Zika viruses for host interacting proteins potentially related to the recent epidemic

**DOI:** 10.1101/529370

**Authors:** P.A. Desingu

## Abstract

Zika virus (ZIKV), circulating in more than 70 countries since 2014, is causing severe developmental abnormality to compare to pre-epidemic infection. ZIKV related flaviviruses as the ability to produce subgenomic flaviviral RNAs (sfRNAs) which are associated with pathogenicity in foetal mice. This study, delineate the increased virulence of ZIKV through sfRNA mediated host protein interaction. Phylogenetically ZIKV sfRNA formed three distinct clades of African, Asian and American strains. Compare with pre-epidemic, the epidemic ZIKV sfRNA has genetic, RNA secondary structure and host protein interacting profile diversity. Interestingly this study found that ZIKV sfRNA interacting proteins involved in the neuronal development, differentiation, proliferation, and apoptosis along with spermatogenesis, host immunity and viral pathogenesis. The difference in the interaction profile and interaction strength between pre-epidemic and epidemic virus could be the reason for the increased virulence of the recent epidemic viruses. Targeting this protein will be the potent choice for antiviral drug designing.

## Introduction

Zika virus (ZIKV), a mosquito-borne RNA virus belongs to the family Flaviviridae, genus Flavivirus. ZIKV infection causes a dengue-like mild disease, with exceptional cases of Guillain-Barre syndrome (GBS) and microcephaly between 1947 to 2006 in Africa and Asia (1–4). Intermittent ZIKV outbreaks have been reported from Southeast Asia during the past 12 years without direct evidence of any neurological disorder (5–9). However, since 2015, severe ZIKV outbreaks have been documented from many countries in the Americas with neurological complications (microcephaly in fetus and newborns, and GBS). Some studies were reported on the mutation in the ZIKV which linked with the rapidly expanding epidemic. Other studies were reported that African strain MR766 produces higher viral titers and more apoptosis in cell cultures (2, 10–13). The outcome of flaviviral infection contributed by many factors among this one of the determinants is subgenomic flaviviral RNAs (sfRNAs).

The single-stranded flaviviral genomic RNA (gRNA) is generally degraded by the host enzyme Xrn1 (5’-3’ exoribonuclease 1). The degradation of the flaviviral gRNA proceeds from 5’ untranslated region (5’UTR), but is halted near the beginning of 3’UTR due to the presence of pseudoknot structure leading to the production of 300 to 500-nucleotide-long subgenomic flaviviral RNAs (sfRNAs) (14, 15). Mutations or deletions within the 3’UTR that inhibit the sfRNA production have been characterized. These mutations significantly attenuate the cytopathic effect and are also less pathogenic when injected to mice. Though transfection of sfRNA without viral components fails to induce significant cell death, it can restore the cytopathacity of a mutant virus which has lost the ability to produce sfRNA (15). ZIKV also produce sfRNA (16, 17), the role of ZIKV sfRNA in the increasing severity of disease outcome is poorly understood. Delineating these mechanisms will be extremely useful to identify novel clues towards determining ZIKV infection. In this study, we performed the comparative analysis of ZIKV sfRNA to identify increase pathogenicity of infection.

## Results

### Phylogenetic analysis of ZIKV sequences

In this study, to find out the genetic diversity of among ZIKV sfRNA, the phylogenetic tree was constructed using the nucleotide sequences retrieved from NCBI public database. This analysis revealed that African, Asian and American sequences were formed in different distinct clades. In the case of Asian strains, three distinct clades were noticed (Figure 1). Clade I is formed by Malaysia/1966 viruses; clade II is formed by Chinese and Thailand sequences; and clade III is French Polynesia, Australian, Japanese and Chinese sequences (Figure 1). In case of African strains, MR766 and DAK strains were formed distinct clades. Interestingly, nucleotide sequences from strain MR766 formed the different distinct clade (Figure 1).

**Figure 1:**
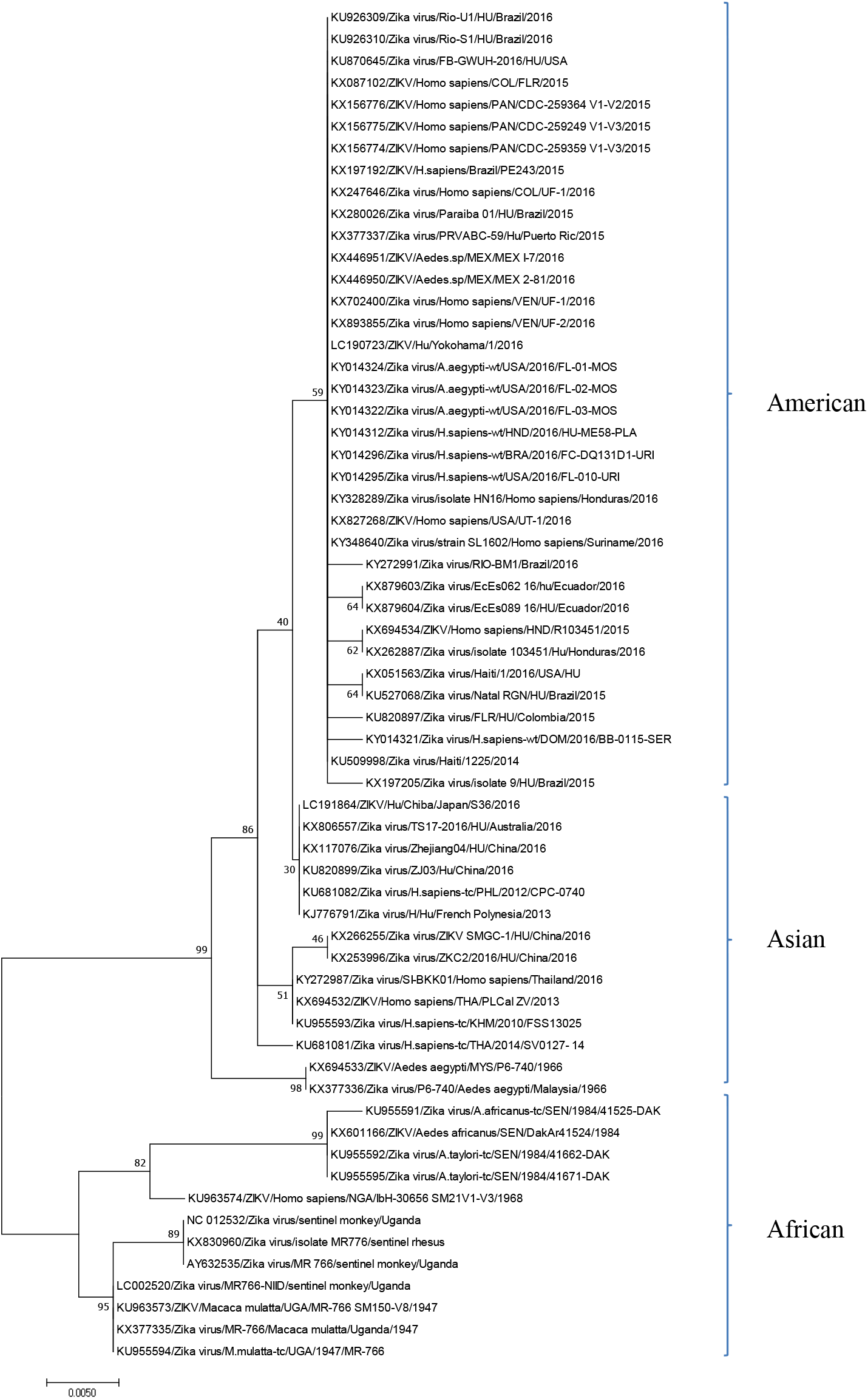
Phylogenetic tree of ZIKV sfRNA sequences. Phylogenetic tree was constructed using the Maximum Likelihood method based on the Kimura 2-parameter model. Initial tree(s) for the heuristic search were obtained automatically by applying Neighbor-Join and BioNJ algorithms to a matrix of pairwise distances estimated using the Maximum Composite Likelihood (MCL) approach, and then selecting the topology with superior log likelihood value. The tree is drawn to scale, with branch lengths measured in the number of substitutions per site. The analysis involved 62 nucleotide sequences. All positions containing gaps and missing data were eliminated. There were a total of 414 positions in the final dataset. Evolutionary analyses were conducted in MEGA7.

### ZIKV sfRNA genetic diversity among African strains

The MR766 sequences, NC_012532/Zika virus/sentinel monkey/Uganda, KX830960/Zika virus/isolate MR776/sentinel rhesus and AY632535/Zika virus/MR 766/sentinel monkey/Uganda sequences formed clade I (Figure 1). NC_012532/Zika virus/sentinel monkey/Uganda and AY632535/Zika virus/MR 766/sentinel monkey/Uganda sequences had 100% identity with 428bp length. KX830960/Zika virus/isolate MR776/sentinel rhesus virus has 429 bp lengths and it has extra nucleotide ‘C’ at the position of 332 (Supplementary Figure 1). The LC002520/Zika virus/MR766-NIID/sentinel monkey/Uganda, KU963573/ZIKV/Macaca mulatta/UGA/MR-766_SM150-V8/1947, KX377335/Zika virus/MR-766/Macaca mulatta/Uganda/1947 and KU955594/Zika virus/M.mulatta-tc/UGA/1947/MR-766 sequences formed clade II (Figure 1) with 100% identity of 429 bp lengths among these sequences. Compared with clade I, the clade II sequences have extra nucleotide ‘C’ at the position of 332 and mutations at T396A and T397G (Supplementary Figure 1). These results revealed that MR766 may be mutated in the laboratory condition at different passage. In comparison with MR766, DAK sequences have the mutation at C4T, T11C, G13A, G112T, G130A, C234T, C257T, G279A, T425G, T427C and C428T (Supplementary Figure 2).

### Comparison of ZIKV sfRNA secondary structure among African strains

This study next tested whether the mutations in the African strains could lead to change in the sfRNA secondary structure. The (i) Minimum Free Energy (MFE) structure drawing encoding base-pair probabilities and positional entropy; (ii) Centroid structure drawing encoding base-pair probabilities and positional entropy; and (iii) Mountain plot representation of the MFE, the thermodynamic ensemble, and the centroid structure revealed that MR766 reference sequence (AY632535), MR766 sequence which has extra nucleotide ‘C’ at the position of 332 (KX830960), clade II MR766 sequences which have extra nucleotide ‘C’ at the position of 332 and mutations of T396A and T397G (KU955594) and DAK strain (KU955595) has significant sfRNA structural diversity (Figure 2). The sfRNA secondary structure diversity may influence the pathogenicity of ZIKV.

**Figure 2:**
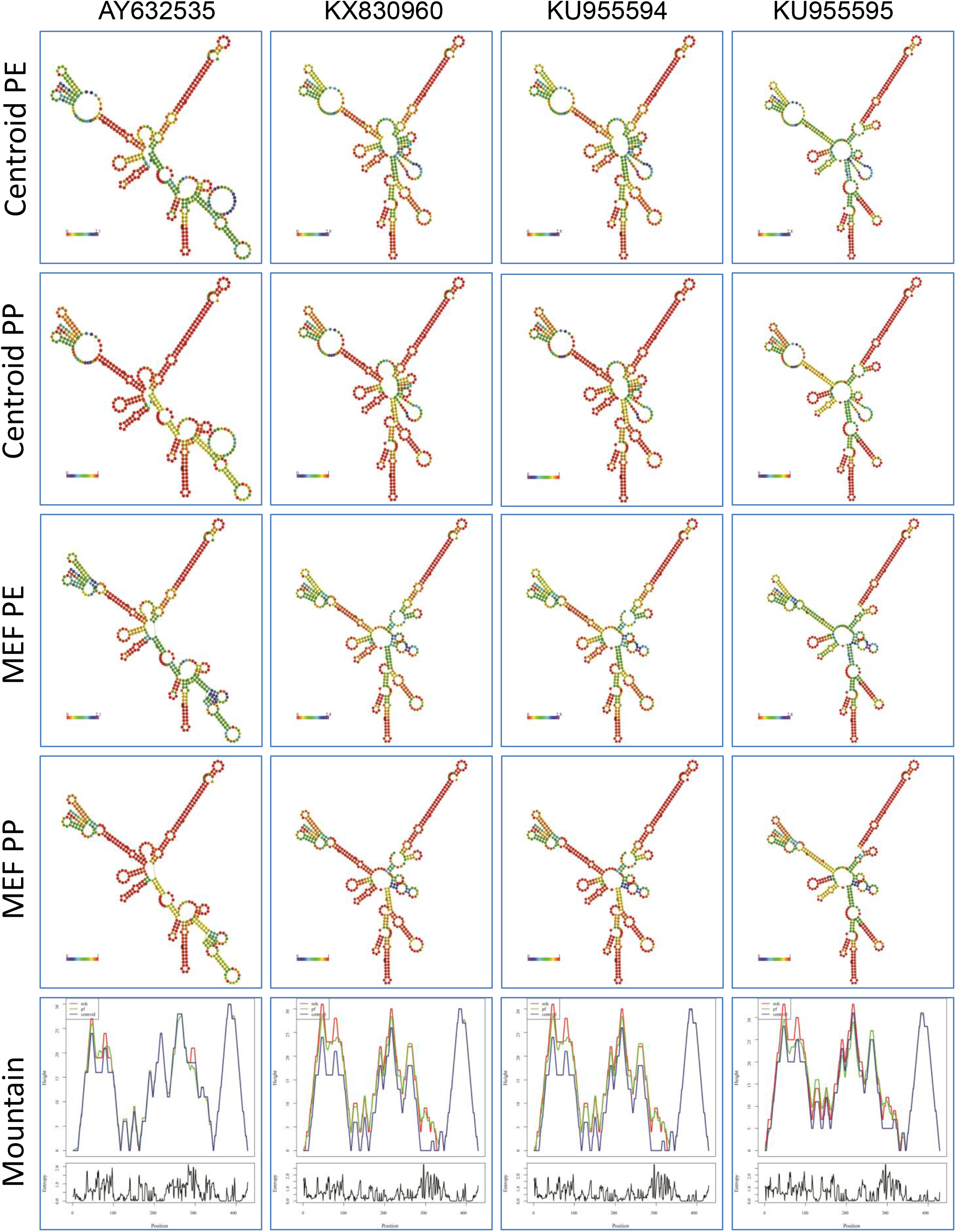
Comparison of ZIKV sfRNA secondary structure among African strains. MR766 reference sequence (AY632535), MR766 sequence which has extra nucleotide ‘C’ at the position of 332 (KX830960), clade II MR766 sequences which have extra nucleotide ‘C’ at the position of 332 and mutations of T396A and T397G (KU955594) and DAK strain (KU955595). Centroid PE-Centroid structure drawing encoding positional entropy; Centroid PP-Centroid structure drawing encoding base-pair probabilities; MEF PE-Minimum Free Energy (MFE) structure drawing encoding positional entropy; MEF PP-Minimum Free Energy (MFE) structure drawing encoding base-pair probabilities; and Mountain-Mountain plot representation of the MFE, the thermodynamic ensemble, and the centroid structure.

### Comparison of host proteins interaction with ZIKV sfRNA among African strains

The contradicting reports are available about the African ZIKV (MR766) virulence in comparison to recent epidemic ZIKV (2, 10–13). This study thought that the controversial results of MR766 virus pathogenicity could be due to the accumulation of mutation and change in the secondary structure in the sfRNA of the MR766 virus at different passage. To explore the sfRNA mediated pathogenicity, we planned to predict the ZIKV sfRNA interacting host proteins. The ZIKV sfRNA interacting proteins were identified using online software (http://s.tartaglialab.com/page/catrapid_omics_group) and top 120 interacting proteins were used in the study. Interestingly, ZIKV sfRNA interact with host proteins involves in the neuronal development, differentiation and death, embryonic development, reproduction, cell division, immunity and virus pathogenesis associated proteins. The details are presented as follows, MR766 reference sequence (AY632535), MR766 mutants (KX830960 and KU955594) and DAK (KU955595).

#### (i) The host proteins involved in the neuronal development, differentiation and death

The host proteins involved in the brain development which interact with ZIKV sfRNA are PROX1, NOVA1, TBR1, ARNT2, PRDM8, SATB2, ZNF131, ZBTB18, CPEB1, FOXP2, FMR1 and GRHL2 (Figure 3a-c). Among these PROX1 and NOVA1 interact with all four sequences of African strain. The TBR1 and ARNT2 interact only with strain MR766; the FMR1and GRHL2 interacts only with Dak strain. PRDM8 interact only with mutant strains of MR766 and Dak. SATB2 specifically interact with mutant strains of MR766. Reference MR766 strain specifically interacts with ZBTB18 and CPEB1. Dak strain specifically interacts with FMR1 and GRHL2. The host proteins involved in the neuronal differentiation TCF4, TCF3, POU6F2 and TCF12 are interacting with ZIKV sfRNA; in addition to FOXO3, CREB3L2 and MEF2D have a role in neuronal death (Figure 3a-c). The TCF3 and FOXO3 protein interact with all four sequences of African strains. TCF4 and TCF12 interact only with mutant strains of MR766 and Dak. Reference MR766 strain specifically interacts with CREB3L2 and MEF2D. In addition, The HNRNPD and ENOX1 have a role in biological clock were interact with ZIKV sfRNA (Figure 3a-c). The ENOX1 protein interacts with all four sequences of African strains. The HNRNPD interact only with mutant strains of MR766 and Dak. Further, SRSF10, C3H14 and PHF21A proteins which regulate RNA biogenesis in neurons were interacted with ZIKV sfRNA. SRSF10 and PHF21A protein interact with all four sequences of African strains. C3H14 interact only with mutant strains of MR766 and Dak. The overall Z score, Discriminative Power (%) [DP%] and Interaction Strength (%) [IS%] of these interacting proteins revealed that the mutant strains of MR766 have more interaction score compared to reference MR766 and Dak strain has least interaction score (Figure 3a-c). These results revealed that mutation in the MR766 strain which accumulates in laboratory passages might be increased the neuronal pathogenicity of the virus.

**Figure 3:**
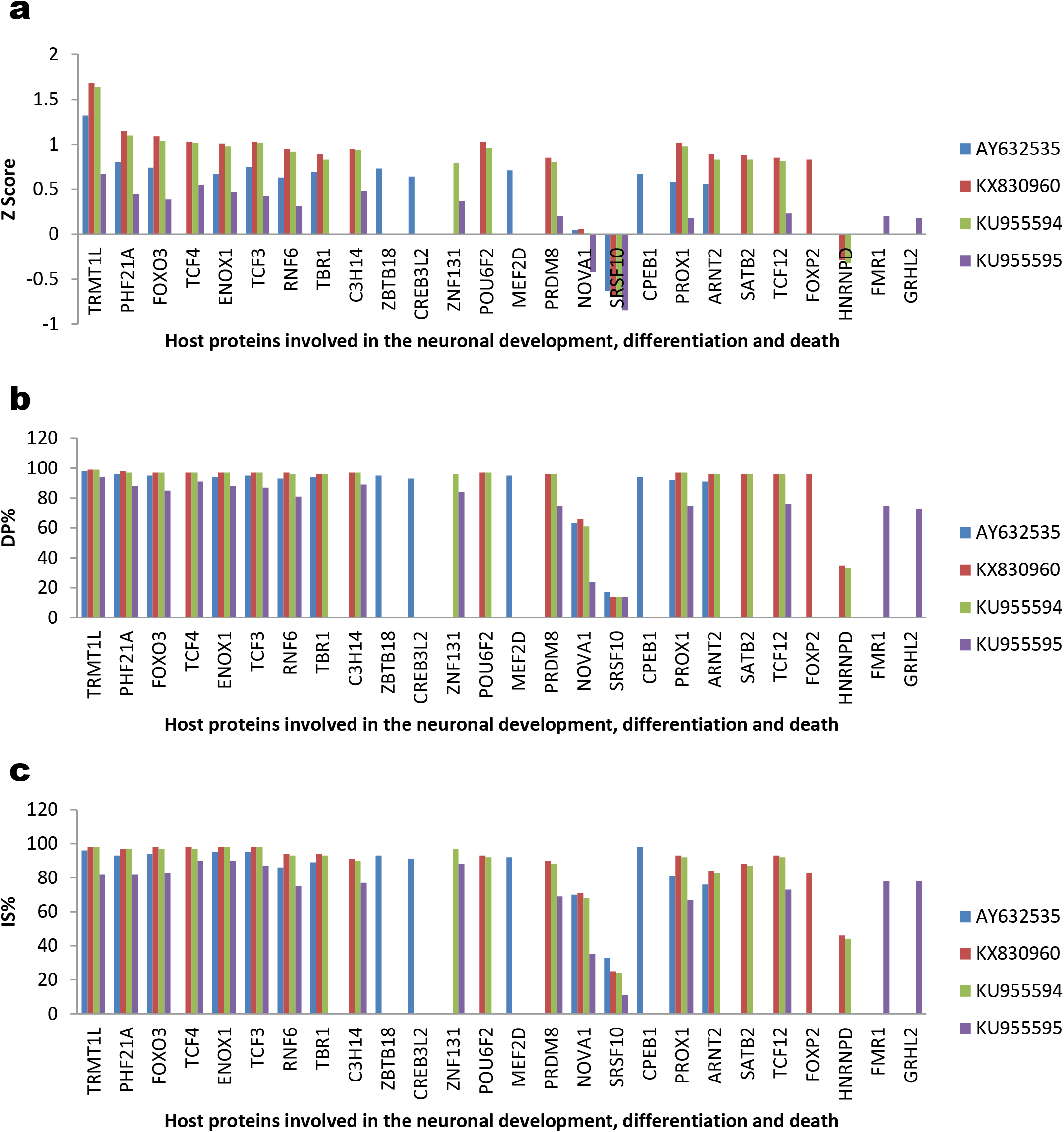
Comparison of the interaction of host proteins involved in the neuronal development, differentiation and death with ZIKV sfRNA among African strains. MR766 reference sequence (AY632535), MR766 sequence which has extra nucleotide ‘C’ at the position of 332 (KX830960), clade II MR766 sequences which have extra nucleotide ‘C’ at the position of 332 and mutations of T396A and T397G (KU955594) and DAK strain (KU955595). a) Z score; b) DP%; and c) IS%

#### (ii) The proteins involved in the embryonic development

The host proteins involved in the embryonic developmental interact with ZIKV sfRNA virus are SCML2, TBX2, PROX1, RTF1, ELAVL1, ATF6, SETDB2, PUM3 and ZBTB16 (Figure 4a-c). SCML2, TBX2, PROX1, RTF1, ELAVL1, ATF6 and SETDB2 protein interact with all the four sequences of African strains. PUM3 interact only with mutant strains of MR766 and Dak. ZBTB16 specifically interact with Dak strain. The Z score, DP% and IS% of the mutant strains of MR766 have higher interaction score compared to reference MR766; Dak strain has lesser interaction score compared to reference MR766 (Figure 4a-c). These results suggest that reference MR766 and Dak strain may cause the mild effect on embryonic to development compare to mutant MR766 strains.

**Figure 4:**
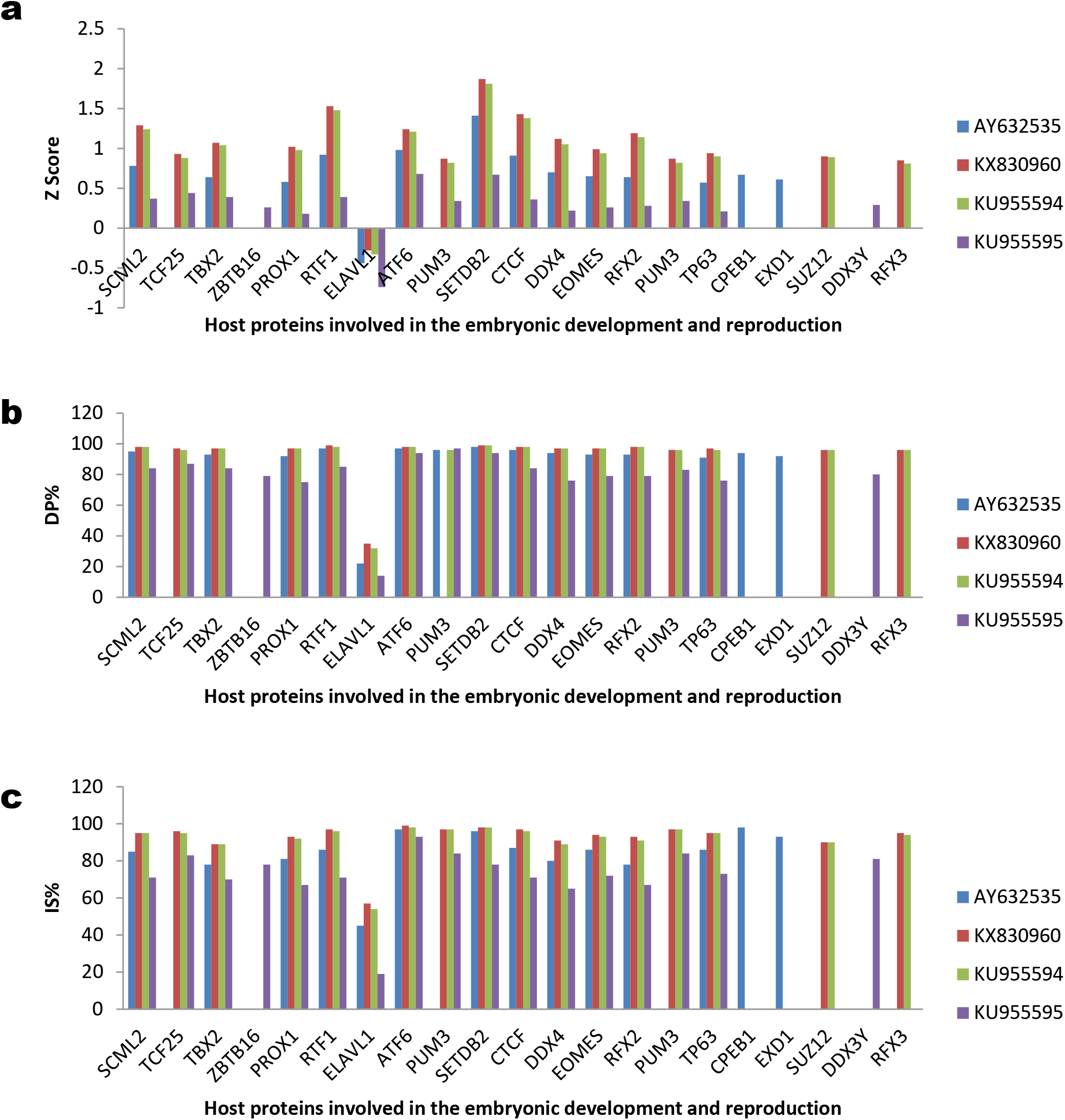
Comparison of the interaction of host proteins involved in the embryonic development and reproduction with ZIKV sfRNA among African strains. MR766 reference sequence (AY632535), MR766 sequence which has extra nucleotide ‘C’ at the position of 332 (KX830960), clade II MR766 sequences which have extra nucleotide ‘C’ at the position of 332 and mutations of T396A and T397G (KU955594) and DAK strain (KU955595). a) Z score; b) DP%; and c) IS%

#### (iii) The proteins involved in the reproduction

Host proteins involved in the spermatogenesis DDX4, RFX2, EXD1, RFX3 and DDX3Y are interacting with ZIKV sfRNA (Figure 4a-c). DDX4 and RFX2 protein interact with all the four sequences of African strains. RFX3 is specifically interacting with mutant strains of MR766. EXD1 and DDX3Y specifically interact with the reference strain of MR766 and Dak respectively. Proteins involved in the female reproduction CTCF, EOMES, PUM3, TP63 and SUZ12 are interacting with ZIKV sfRNA. The CTCF, EOMES and TP63 proteins interact with all the four sequences of African strains. PUM3 interact only with mutant strains of MR766 and Dak. SUZ12 is specifically interacting with mutant strains of MR766. The interaction strength of mutant MR766 is higher than reference MR766 and DAK strains (Figure 4a-c).

#### (iv) The proteins involved in the cell division, differentiation, proliferation and death

The proteins involved in the cell division, differentiation, proliferation and death which interact with ZIKV sfRNA are MYBL2, HIC1, FOXO3, ZGPAT, HDGFL2, DEAF1, POLH, DBF4, PIF1, FBXW7, ZBTB48, MITF, PTBP1, HNRNPK, DBF4B, AGGF1, SKI, ZNF16, FOXK1, ZNF35, SP6 and SARNP (Figure 5a-c). The MYBL2, HIC1, FOXO3, HDGFL2, POLH, DBF4, PIF1, FBXW7, PTBP1, HNRNPK, MRE11, KIF2C and TMPO protein interact with all four sequences of African strains. FBXW7 is interacting with reference and mutant strains of MR766. ZBTB48 is interacting only with mutant strains of MR766 and Dak. SKI, ZNF16 and AKAP8 are interacting only with mutant strains of MR766. The ZGPAT, MITF and SUMO1 are specifically interacting with the reference sequence of MR766. These proteins have shown the same trend that mutant MR766 possess higher interaction ability (Figure 5a-c).

**Figure 5:**
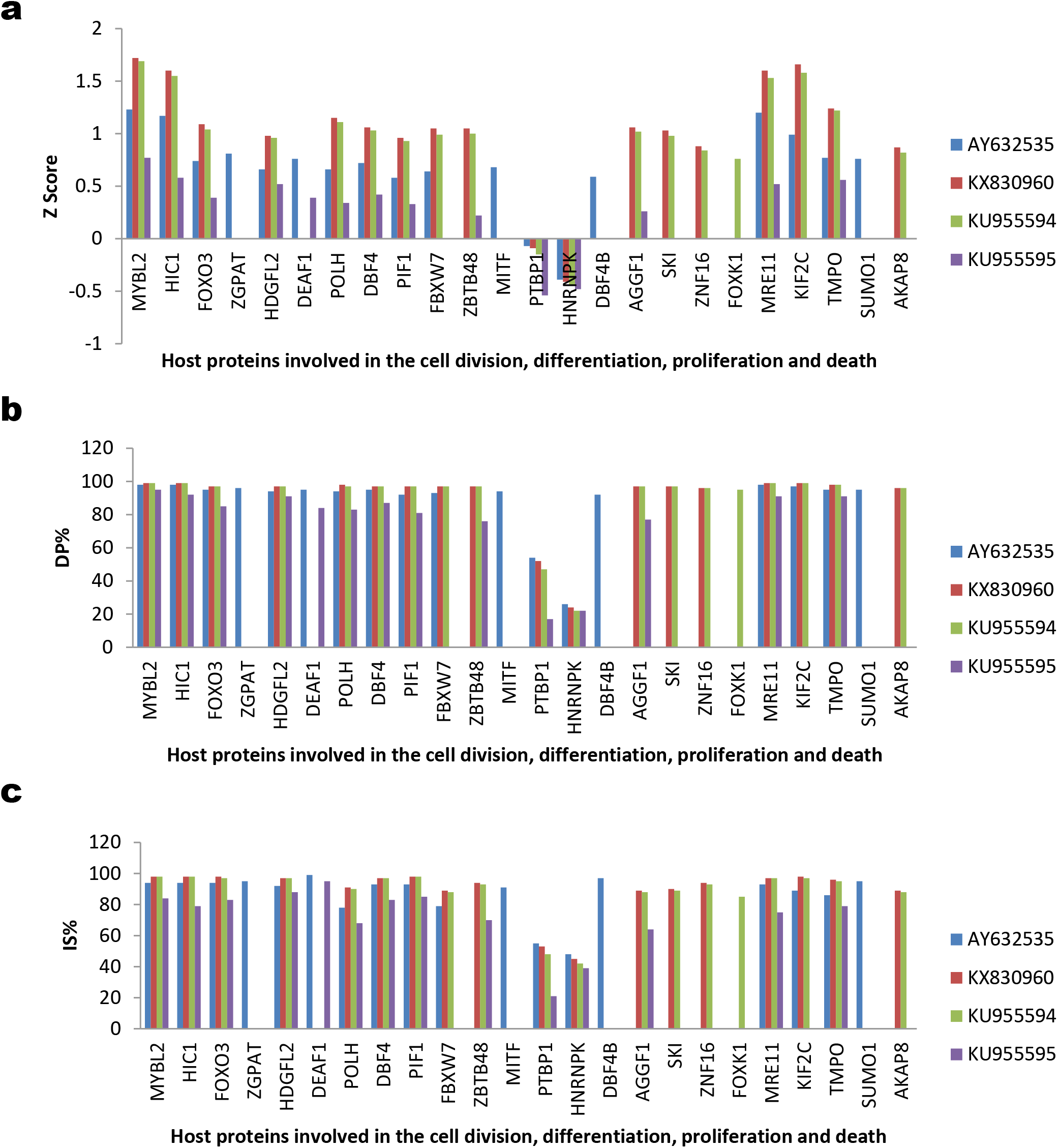
Comparison of the interaction of host proteins involved in the cell division, differentiation, proliferation and death with ZIKV sfRNA among African strains. MR766 reference sequence (AY632535), MR766 sequence which has extra nucleotide ‘C’ at the position of 332 (KX830960), clade II MR766 sequences which have extra nucleotide ‘C’ at the position of 332 and mutations of T396A and T397G (KU955594) and DAK strain (KU955595). a) Z score; b) DP%; and c) IS%

#### (v) The proteins involved in the host immunity

Proteins involved in the host immunity ZBTB1, NKRF, SFPQ, CALCOCO1, TMPO, BANP, BCL6, RBM14, NFE2L3 and ZBTB20 are interacting with ZIKV sfRNA (Figure 6a-c). The ZBTB1, NKRF, SFPQ, CALCOCO1 and TMPO proteins interact with all the four sequences of African strains. BANP and BCL6 proteins are interacting with reference and mutant strains of MR766. ZBTB20 is interacting with mutant strains of MR766 and Dak. NFE2L3 is interacting with mutant strains of MR766. RBM14 is specifically interacting with Dak strain. The Z score, DP% and IS% revealed the similar trend that mutant MR766 has more interaction strength (Figure 6a-c).

**Figure 6:**
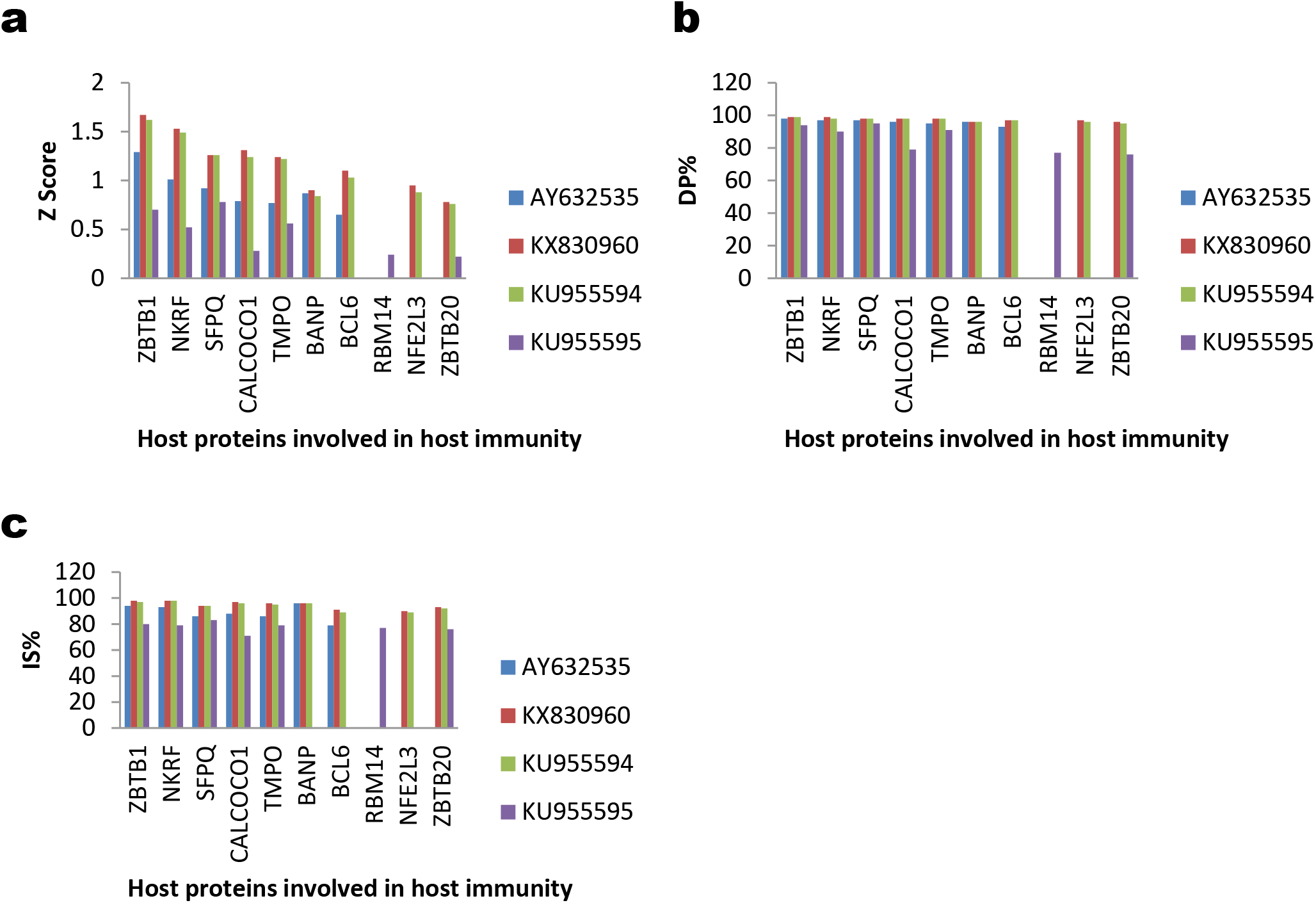
Comparison of the interaction of host proteins involved in host immunity with ZIKV sfRNA among African strains. MR766 reference sequence (AY632535), MR766 sequence which has extra nucleotide ‘C’ at the position of 332 (KX830960), clade II MR766 sequences which have extra nucleotide ‘C’ at the position of 332 and mutations of T396A and T397G (KU955594) and DAK strain (KU955595). a) Z score; b) DP%; and c) IS%

#### (vi) The proteins involved in the virus pathogenesis

The proteins involved in the virus pathogenesis which interact ZIKV sfRNA virus are DHX58, TRIM25, REXO1L1P, RTF1, OAS2, CPEB3, CCNT1, RBM14, PCBP1, PCBP2, HNRNPA1, DDX1, RNASEL, PABPC1 and FMR1 (Figure 7a-c). The DHX58, TRIM25, REXO1L1P, RTF1, OAS2, CPEB3, CCNT1, RBM14, PCBP1, PCBP2 and HNRNPA1 protein interacts with all the four sequences of African strains. DDX1 is interacting with mutant strains of MR766 and Dak. RNASEL is interacting with mutant strains of MR766. PABPC1 and FMR1 are specifically interacting with Dak strain. The overall Z score, DP% and IS% of these interacting proteins revealed that the mutant strains of MR766 have more interaction score compared to reference MR766 and Dak strain has least interaction score (Figure 7a-c).

**Figure 7:**
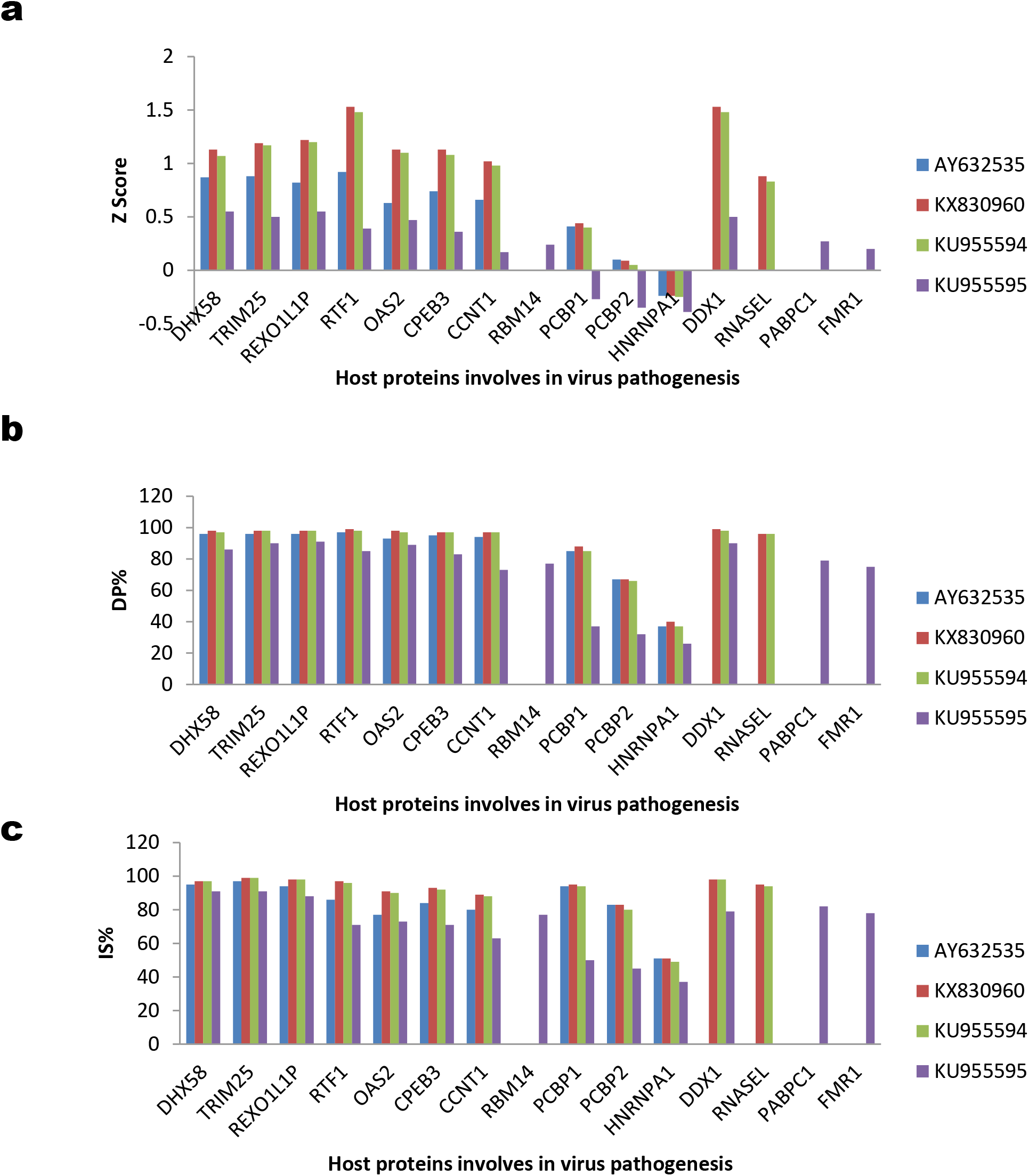
Comparison of the interaction of host proteins involved in the virus pathogenesis with ZIKV sfRNA among African strains. MR766 reference sequence (AY632535), MR766 sequence which has extra nucleotide ‘C’ at the position of 332 (KX830960), clade II MR766 sequences which have extra nucleotide ‘C’ at the position of 332 and mutations of T396A and T397G (KU955594) and DAK strain (KU955595). a) Z score; b) DP%; and c) IS%

### Comparison of sfRNA secondary structure in the genome of Asian and American Zika virus strains

This study next tested whether the genetic diversity in the Asian and American strains could lead to change in the sfRNA secondary structure. The (i) MFE structure drawing encoding base-pair probabilities and positional entropy; (ii) Centroid structure drawing encoding base-pair probabilities and positional entropy; and (iii) Mountain plot representation of the MFE, the thermodynamic ensemble, and the centroid structure revealed that three different clades of Asian strains and American (428 and 429 bp size sfRNA) strains shown significant sfRNA structural diversity at Centroid structure; while MEF structure, only clade I of Asian strain has significant diversity (Figure 8).

**Figure 8:**
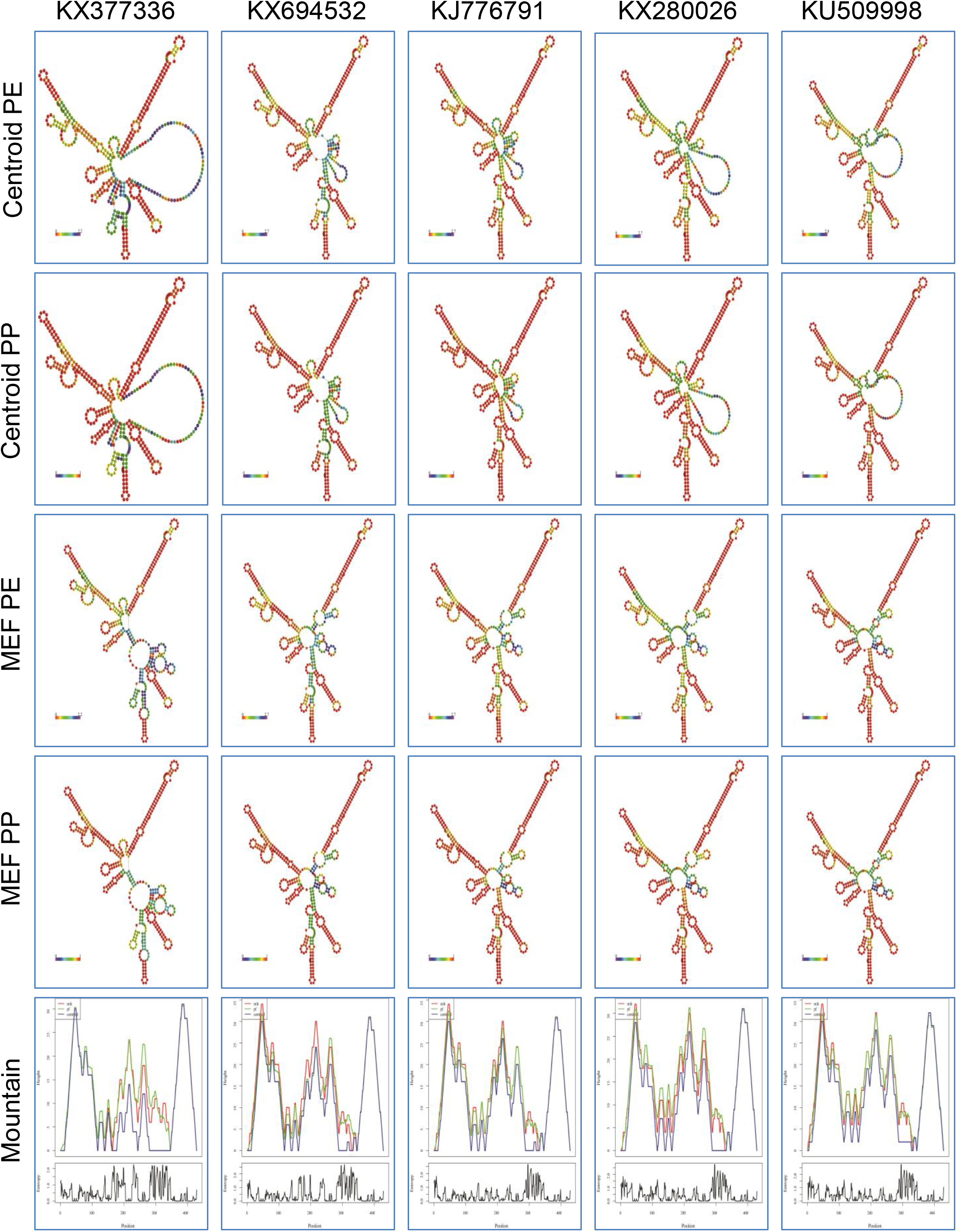
Comparison of ZIKV sfRNA secondary structure of pre-epidemic and epidemic ZIKV. The details are presented as follows, African strains [MR766 (AY632535), DAK (KU955595)], Asian strains [clade I (Malaysia/1966-KX377336); clade II (Thailand-KX694532); and clade III (French Polynesia-KJ776791)] and American strains [Strain with 428 bp size sfRNA (Brazil/2015-KX280026) and Strain with 429 bp size sfRNA (Haiti/2014-KU509998)]. Centroid PE-Centroid structure drawing encoding positional entropy; Centroid PP-Centroid structure drawing encoding base-pair probabilities; MEF PE-Minimum Free Energy (MFE) structure drawing encoding positional entropy; MEF PP-Minimum Free Energy (MFE) structure drawing encoding base-pair probabilities; and Mountain-Mountain plot representation of the MFE, the thermodynamic ensemble, and the centroid structure.

### Comparison of host proteins interacting with sfRNA in the genome of pre-epidemic and epidemic Zika virus strains

The limited direct evidence is available about Asian strains of ZIKV mediated neurological disorder (5–9). In the phylogenetic analysis, sfRNA revealed that Asian strains are out branched with American strains of ZIKV. Asian strains of ZIKV have formed three different distinct clades. In addition, American strains of ZIKV have two different 428 bp and 429 bp size of sfRNA. Further, genetic diversity and secondary structural diversity among pre-epidemic and epidemic Zika virus sfRNA might be leads to increased pathogenicity. To explore the sfRNA mediated pathogenicity, we planned to predict the ZIKV sfRNA interacting host proteins. The details are presented as follows, African strains [MR766 (AY632535), DAK (KU955595)], Asian strains [clade I (Malaysia/1966-KX3773360); clade II (Thailand-KX694532); and clade III (French Polynesia-KJ776791)] and American strains [Strain with 428 bp size sfRNA (Brazil/2015-KX280026) and Strain with 429 bp size sfRNA (Haiti/2014-KU509998)].

#### (i) The proteins involved in the neuronal developmental, differentiation and death

The ZIKV sfRNA interacting host proteins involved in brain development were compared (Figure 9a-c). NOVA1 is interacting with all seven ZIKV sequence analysed. ZBTB18 is with all sequence except DAK strain. The TBR1 is interacting with MR766, Malaysia/1966 and American strains. PRDM8 is interacting with DAK, Malaysia/1966 and American strains. ZNF131 is interacting with DAK and American strains. PROX1 is interacting with African strains and American strain Haiti/2014. CPEB1 is specifically interacting with MR766 and Malaysia/1966. ARNT2 is specifically interacting with MR766 and American strain Haiti/2014. FMR1 and GRHL2 are specific to DAK strain. ATXN1L is specific to American strain Haiti/2014. Then, the ZIKV sfRNA interacting host proteins involved in neuronal differentiation were compared (Figure 9a-c). The TCF3 is interacting with African strains, Malaysia/1966 and American strains. TCF4 is specific to DAK and American strains. POU6F2 is specific to American strains. TCF12 and BEX1 are specifically interacting with DAK and French Polynesia strains respectively. In case of neuronal death, FOXO3 is interacting with all seven ZIKV sequence analysed; CREB3L2 is with all sequence except DAK strain; MEF2D is interacting with MR766, Asian strains and American strain Brazil/2015 (Figure 9a-c). Protein involves in biological clock ENOX1 is specifically interacting with African and American strains (Figure 9a-c). Further, proteins which regulate RNA biogenesis in neurons were compared. PHF21A is interacting with all seven ZIKV sequence analysed; SRSF10 is with all sequence except French Polynesia strain; C3H14 is specific to DAK and American strains (Figure 9a-c). The overall Z score, DP% and IS% of African strains are lesser than American strains; the Asian strains are unable to interact with some common host proteins interacting with African and American strains (Figure 9a-c). These results suggest that the loss of interaction with the proteins involved in neuronal developmental, differentiation and death by Asian strains leads to less neuronal damage by these strain. Further, lesser interaction strength of African strains compares to American strains may lead to severe neuronal damage in the recent epidemic of ZIKV infection.

**Figure 9:**
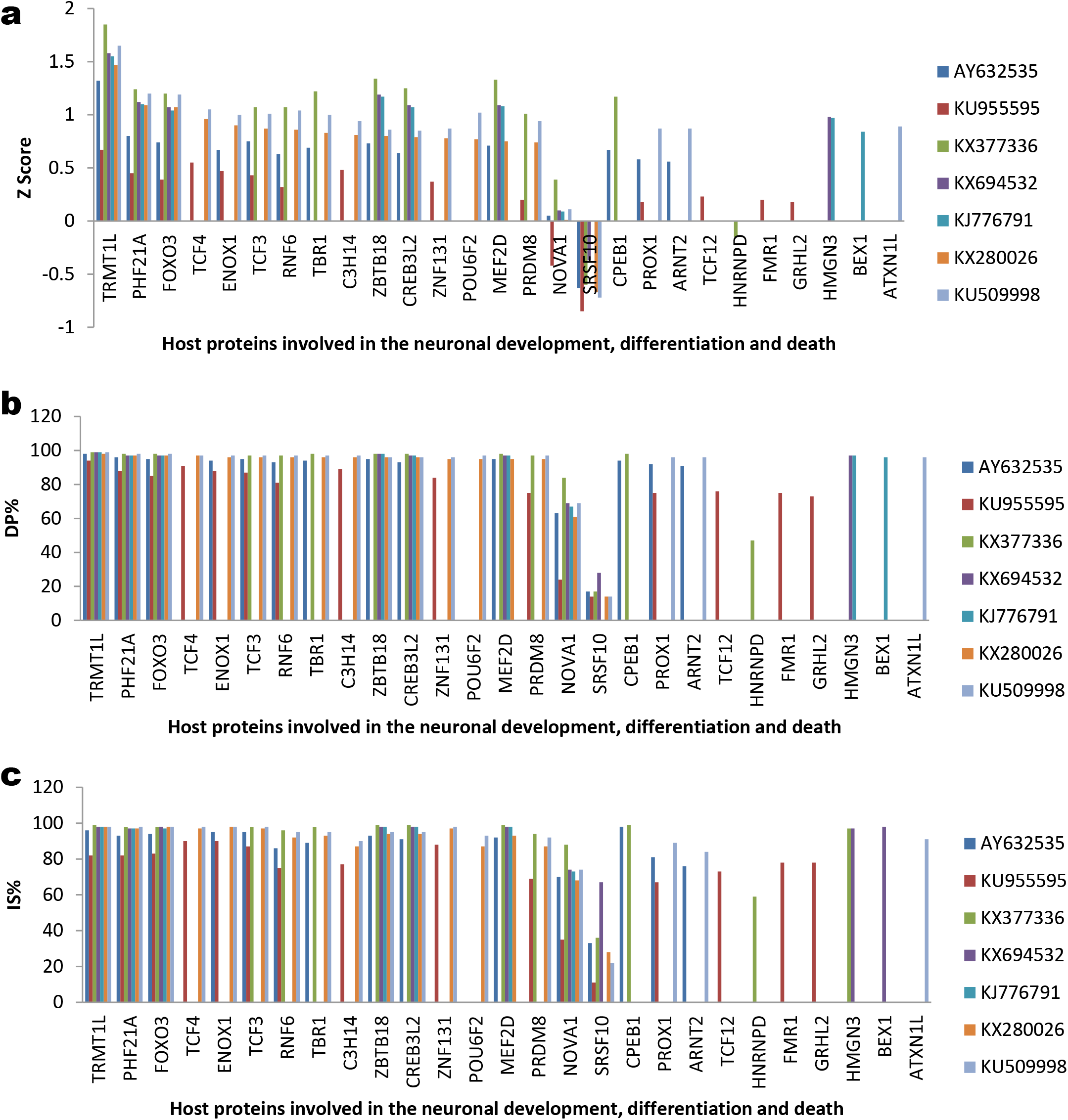
Comparison of pre-epidemic and epidemic ZIKV sfRNA interacting host proteins involved in the neuronal development, differentiation and death. The details are presented as follows, African strains [MR766 (AY632535), DAK (KU955595)], Asian strains [clade I (Malaysia/1966-KX377336); clade II (Thailand-KX694532); and clade III (French Polynesia-KJ776791)] and American strains [Strain with 428 bp size sfRNA (Brazil/2015-KX280026) and Strain with 429 bp size sfRNA (Haiti/2014-KU509998)]. a) Z score; b) DP%; and c) IS%

#### (ii) The proteins involved in the embryonic development

The ZIKV sfRNA interacting host proteins involved in embryonic development were compared (Figure 10a-c). RTF1, ELAVL1, ATF6 and SETDB2 are interacting with all seven ZIKV sequence analysed. SCML2 and TBX2 are interacting with African strains, Malaysia/1966 and American strains. TCF25 and PUM3 are interacting with DAk and American strains. HOPX, HMX1 and HMGN3 are specific to Thailand and French Polynesia strains. PROX1 is specific to African and American strain Haiti/2014. ZBTB16 is specific to DAK and American strain Brazil/2015. The interaction strength lesser for African strains and Asian strain lost ability to interact with common interacting proteins of African and American Strains (Figure 10a-c).

**Figure 10:**
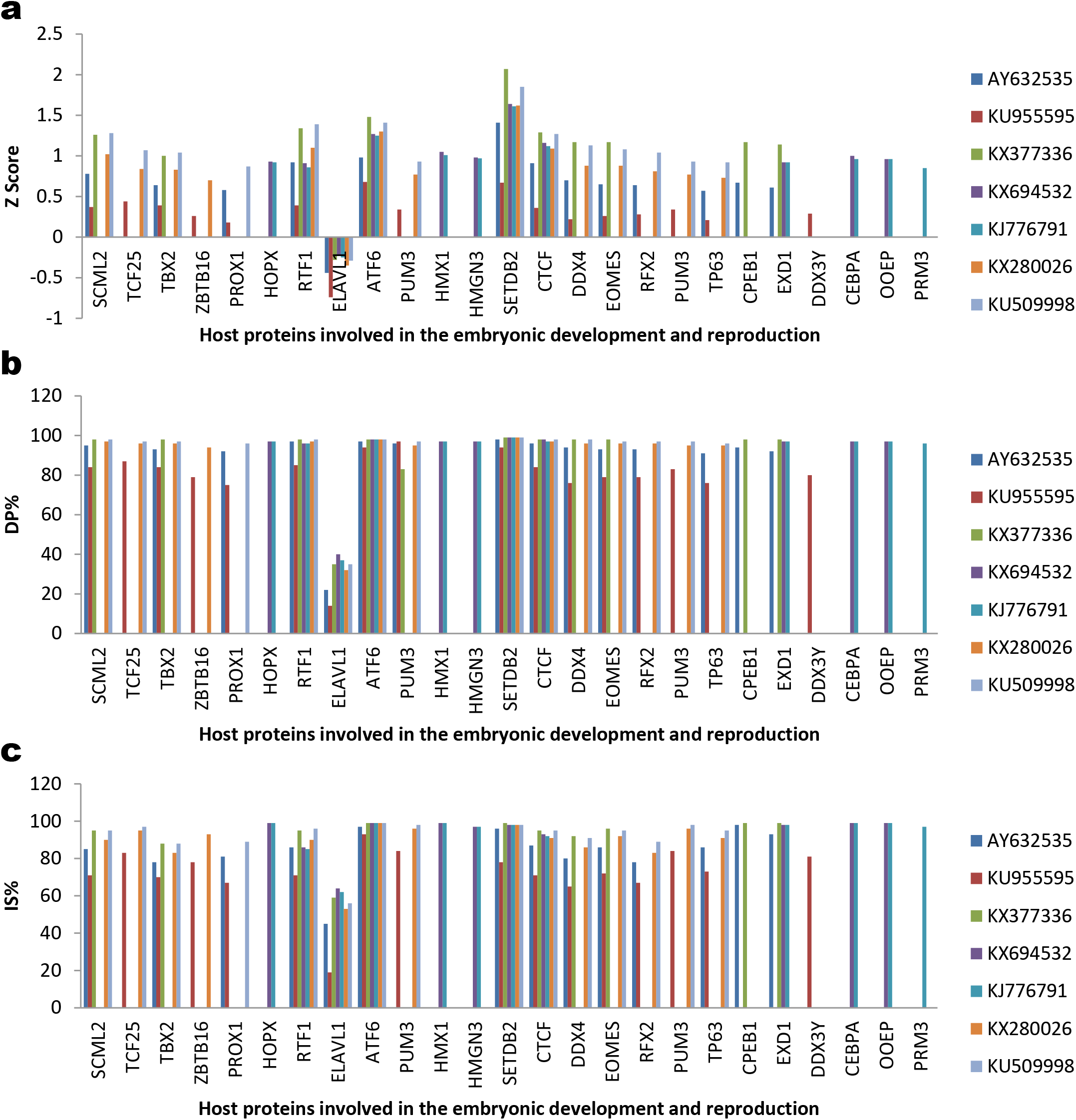
Comparison of pre-epidemic and epidemic ZIKV sfRNA interacting host proteins involved in the embryonic development and reproduction. The details are presented as follows, African strains [MR766 (AY632535), DAK (KU955595)], Asian strains [clade I (Malaysia/1966-KX377336); clade II (Thailand-KX694532); and clade III (French Polynesia-KJ776791)] and American strains [Strain with 428 bp size sfRNA (Brazil/2015-KX280026) and Strain with 429 bp size sfRNA (Haiti/2014-KU509998)]. a) Z score; b) DP%; and c) IS%

#### (iii) The proteins involved in the reproduction

The ZIKV sfRNA interacting host proteins involved in the spermatogenesis were compared (Figure 10a-c). DDX4 is specific to African strains, Malaysia/1966 and African strains; RFX2 is specific to African strains and American strains; EXD1 is specific to MR766 and Asian strains; DDX3Y is specific to DAK strain; PRM3 is specific to French Polynesia strain. The ZIKV sfRNA interacting host proteins involved in the female reproduction were compared. CTCF is interacting with all seven ZIKV sequence analysed. EOMES is specific to African strains, Malaysia/1966 and African strains. The TP63 is interacting with African strains and African strains. PUM3 is interacting with DAk and American strains. CPEB1 is specific to MR766 and Malaysia/1966. CEBPA and OOEP are specifically interacting with Thailand and French Polynesia strains (Figure 10a-c). These results suggest that African strains may cause the mild effect on reproduction to compare to American strains.

#### (iv) The proteins involved in the cell division, differentiation, proliferation and death

The proteins involved in the cell division, differentiation, proliferation and death which interact with ZIKV sfRNA virus were compared (Figure 11a-c). HIC1, FOXO3, DEAF1, PTBP1, HNRNPK and MRE11 are interacting with all seven ZIKV sequence analysed. MYBL2 is interacting with all strains except Thailand strain. MITF, KIF2C and SUMO1 are interacting with MR766, Asian strain and American strain Brazil/2015. HDGFL2, POLH, DBF4 and TMPO are interacting with African strains, Malaysia/1966 and American strains.

**Figure 11:**
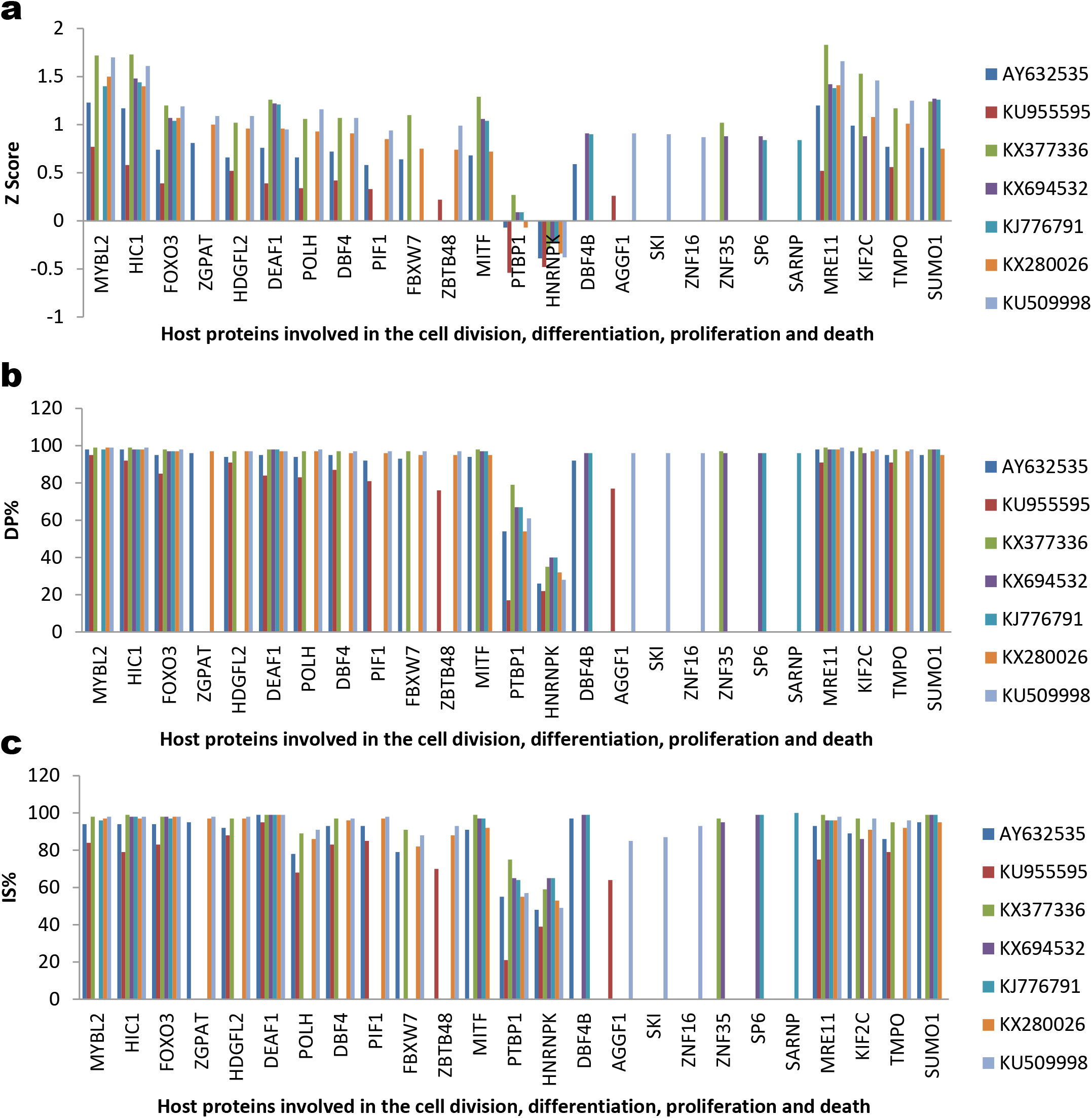
Comparison of pre-epidemic and epidemic ZIKV sfRNA interacting host proteins involved in the cell division, differentiation, proliferation and death. The details are presented as follows, African strains [MR766 (AY632535), DAK (KU955595)], Asian strains [clade I (Malaysia/1966-KX377336); clade II (Thailand-KX694532); and clade III (French Polynesia-KJ776791)] and American strains [Strain with 428 bp size sfRNA (Brazil/2015-KX280026) and Strain with 429 bp size sfRNA (Haiti/2014-KU509998)]. a) Z score; b) DP%; and c) IS%

PIF1 is interacting with African strains and American strains. ZGPAT is specific to MR766 and American strains. ZBTB48 is specific to DAK and American strains; AGGF1 is specific to DAK and American strain Brazil/2015; ZNF35 is specific to Asian strains Malaysia/1966 and Thailand; SP6 is specific to Asian strains Thailand and French Polynesia; SARNP is specific to French Polynesia strain; SKI and ZNF16 are specific to American strain Brazil/2015 (Figure 11a-c). The interaction trends of African, Asian and American are as similar to neuron specific protein interaction.

#### (v) The proteins involved in the host immunity

The ZIKV sfRNA interacting host proteins involved in the host immunity were compared (Figure 12a-c). ZBTB1, NKRF and SFPQ are interacting with all seven ZIKV sequence analysed. BANP is interacting with all sequence except DAK strain. CALCOCO1 and TMPO are interacting with African strains, Malaysia/1966 and African strains. BCL6 is specific to MR766, Malaysia/1966 and American strains. RBM14 is interacting with DAK and American strains. ZBTB20 is specific to DAk; IKZF1 and ARID3A are specific to Malaysia/1966. The largely interaction score of American strains are higher than African strain (Figure 12a-c).

**Figure 12:**
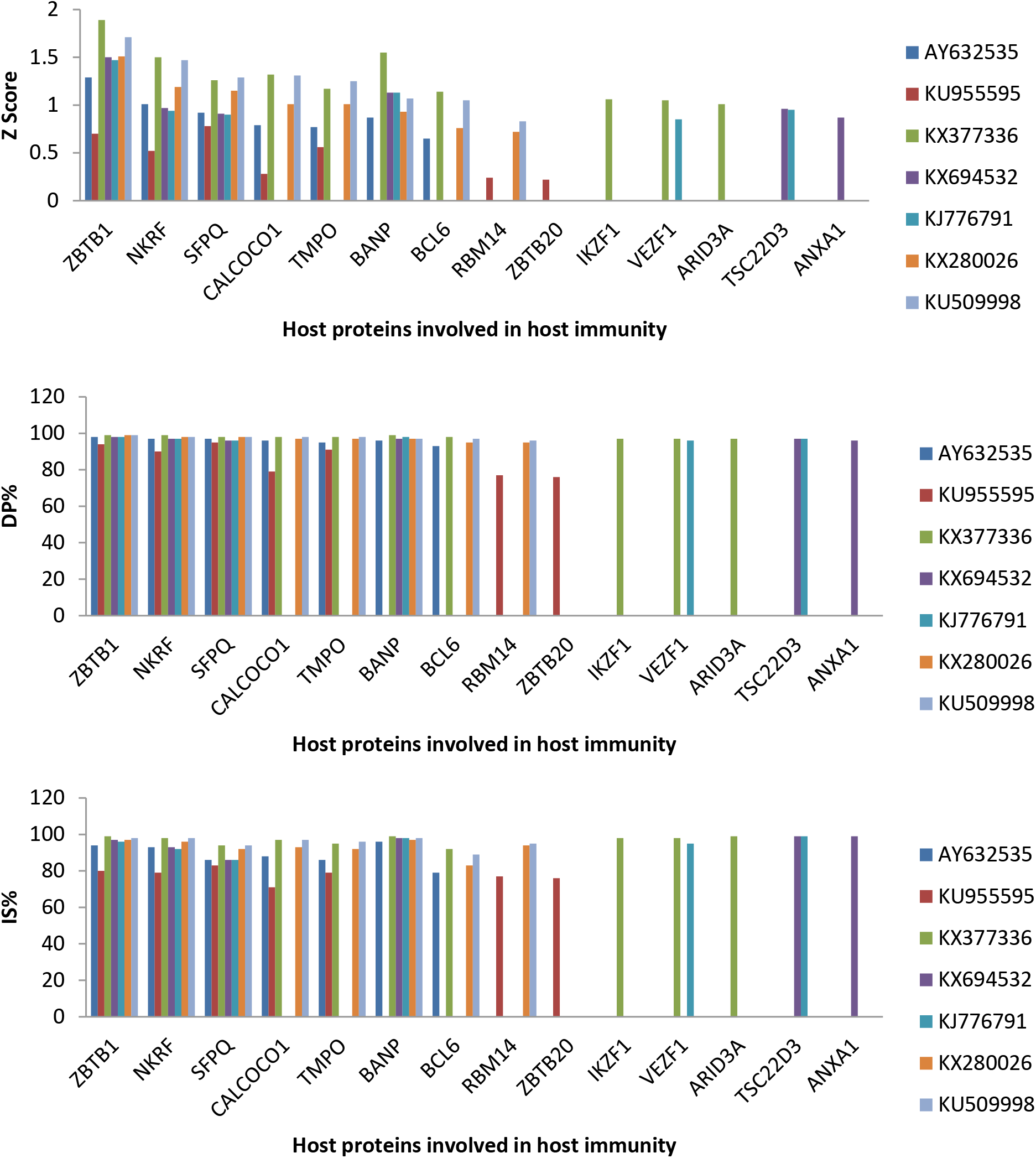
Comparison of pre-epidemic and epidemic ZIKV sfRNA interacting host proteins involved in host immunity. The details are presented as follows, African strains [MR766 (AY632535), DAK (KU955595)], Asian strains [clade I (Malaysia/1966-KX377336); clade II (Thailand-KX694532); and clade III (French Polynesia-KJ776791)] and American strains [Strain with 428 bp size sfRNA (Brazil/2015-KX280026) and Strain with 429 bp size sfRNA (Haiti/2014-KU509998)]. a) Z score; b) DP%; and c) IS%

#### (vi) The proteins involved in the virus pathogenesis

The ZIKV sfRNA interacting host proteins involved in the virus pathogenesis were compared (Figure 13a-c). DHX58, TRIM25, REXO1L1P, RTF1, PCBP1, PCBP2 and HNRNPA1 interacting with all seven ZIKV sequence analysed. CPEB3 and CCNT1 are interacting with African strains, Malaysia/1966 and American strains. DDX1 is interacting with all sequence except MR766 and French Polynesia. OAS2 is specific to African strains and American strains; POLR3C is specific to Asian strains; RBM14 is specific to DAK and American strains. PABPC1 and FMR1 are specific to DAK strain. The general Z score, DP% and IS% of African strains are lesser than American strains (Figure 13a-c).

**Figure 13:**
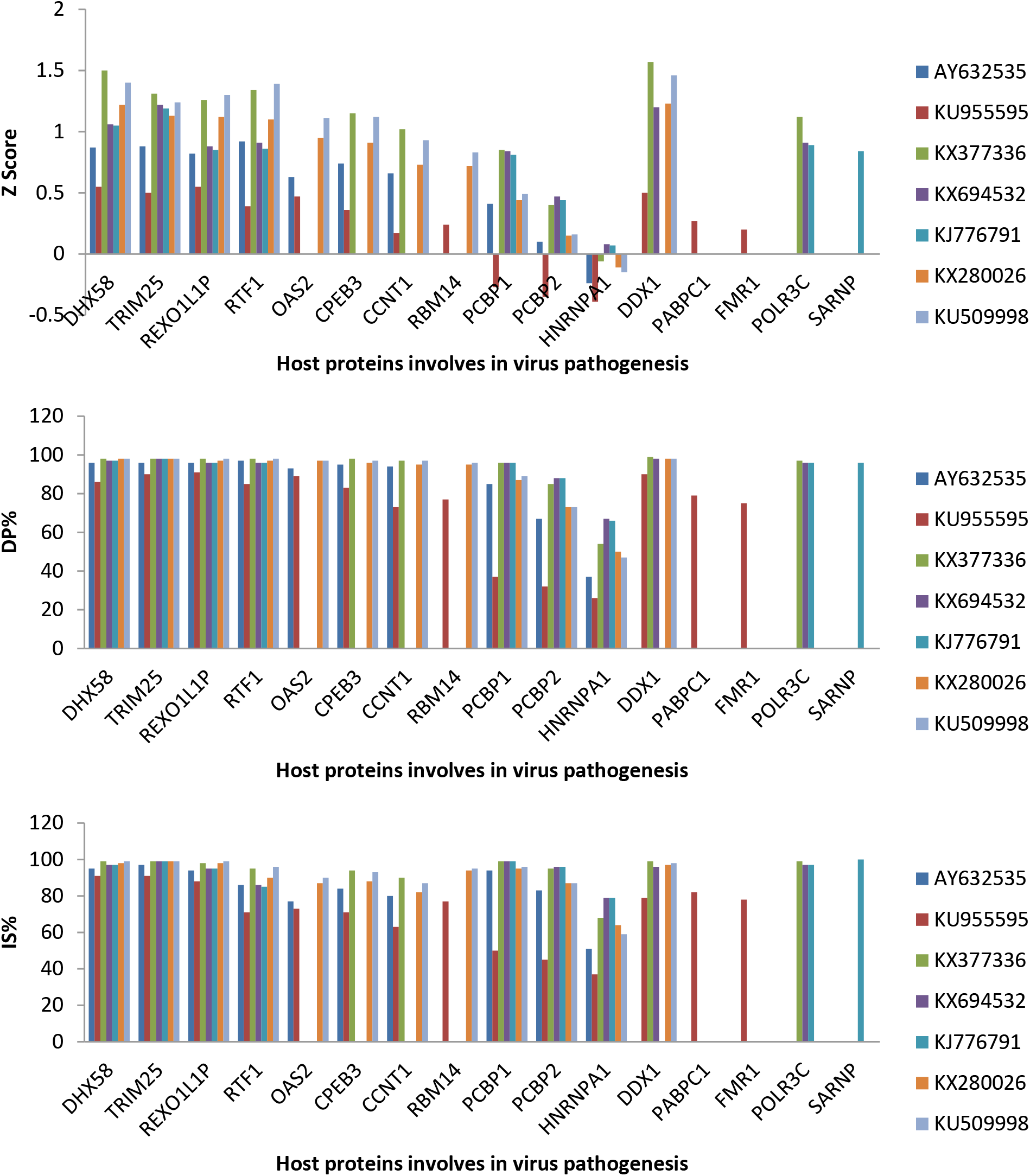
Comparison of pre-epidemic and epidemic ZIKV sfRNA interacting host proteins involved in virus pathogenesis. The details are presented as follows, African strains [MR766 (AY632535), DAK (KU955595)], Asian strains [clade I (Malaysia/1966-KX377336); clade II (Thailand-KX694532); and clade III (French Polynesia-KJ776791)] and American strains [Strain with 428 bp size sfRNA (Brazil/2015-KX280026) and Strain with 429 bp size sfRNA (Haiti/2014-KU509998)]. a) Z score; b) DP%; and c) IS%

### Discussion

The ZIKV causing serious public health emergency in the recent years, it is mandated to identify the underlining mechanism behind the outbreak to design antivirals, diagnostics and vaccine. This study planned to identify the role of ZIKV sfRNA in the recent epidemics of ZIKV mediated developmental abnormality. The flaviviral sfRNA is the product of XRN1 resistant 3’ untranslated region (3’UTR) (14, 15). There are contradicting results are available about the virulence of MR766 strain compare to recent epidemic strains (2, 10-13). This study revealed that the reference MR766 strain has 428 bp length of sfRNA; while the mutant MR766 strains possess 429 bp sizes. The nucleotide length variation could be due to the different passage of MR766 strain in the laboratory as documented earlier (Saiz et al., 2016). The mutant MR766 sfRNA sequence able to form the different secondary structure in MEF and Centroid structure; mutant MR766 sfRNA has the different host protein interacting profile. The overall interaction score of mutant sfRNA MR766 strains is higher than reference MR766. These results suggest that the contradicting results of MR766 strains virulence might be due to testing of the different passage of MR766 virus which has laboratory derived mutations in the sfRNA.

The information about the Asian ZIKV strains related neurological disorders were limited (5–9). In this study, the Asian ZIKV strains were grouped into distinct clades in phylogenetic analysis, possess RNA secondary structure diversity and differ in host protein interacting profile in comparison to American strains. Asian ZIKV sfRNA are interacting with less number of host proteins involves in the neuronal development, differentiation, proliferation and death compare to American strains. These results provide the possible explanation for limited neurological disorders associated with the Asian strains of ZIKV.

Further comparison of African strains (reference MR766 and Dak) and American strains revealed that African strain is totally out branch from American strains of ZIKV sfRNA; distinct sfRNA secondary structure diversity between African and American strains; and overall less Z score, DP% and IS% of interaction score of African sfRNA in comparison to American strain. These results explain the possible reason for the severity of the recent epidemic causing American strains.

The recent epidemic ZIKV causes neurological disorder such as microcephaly is characterized by infecting the developing neurons (2, 18, 19). The mechanism by which ZIKV causing damage in the developing neurons has remains unexplored. This study provided the clue that ZIKV sfRNA could interact with the host proteins involving neurogenesis such as PROX1, NOVA1, TBR1, ARNT2, PRDM8, SATB2, ZNF131, ZBTB18, CPEB1, FOXP2, FMR1 and GRHL2. Other host proteins interacting with ZIKV sfRNA which has role embryonic development are SCML2, TBX2, PROX1, RTF1, ELAVL1, ATF6, SETDB2, PUM3 and ZBTB16. Further, ZIKV affect the differentiation, proliferation and death of neuronal cells (20–24). This study, provided indication proteins has role in the ZIKV pathogenesis in neuronal differentiation; proliferation and death are TCF4, TCF3, POU6F2, TCF12, FOXO3, REB3L2 and MEF2D. Other host protein involves in the cell division differentiation, proliferation and death which interact with ZIKV sfRNA are PROX1, NOVA1, TBR1, ARNT2, PRDM8, SATB2, ZNF131, ZBTB18, CPEB1, FOXP2, FMR1 and GRHL2. ZIKV is reported to cause the reproductive failure such affect spermatogenesis and placenta formation (25–29). The ZIKV sfRNA interacting host proteins involved in spermatogenesis are DDX4, RFX2, EXD1, RFX3 and DDX3Y; and female reproduction are CTCF, EOMES, PUM3, TP63 and SUZ12. The innate immune response, B cell and T cell mediated immune response are reported to have role in the clearance of ZIKV (30–32). The following host immunity related proteins are interacting ZIKV sfRNA: ZBTB1, NKRF, SFPQ, CALCOCO1, TMPO, BANP, BCL6, RBM14, NFE2L3 and ZBTB20. In addition, ZIKV sfRNA interact with the proteins which have role virus pathogenesis are DHX58, TRIM25, REXO1L1P, RTF1, OAS2, CPEB3, CCNT1, RBM14, PCBP1, PCBP2, HNRNPA1, DDX1, RNASEL, PABPC1 and FMR1. Further, identification host protein interacting ZIKV sfRNA sequence and mutating such sequences are needed to understand the role interaction in the viral pathogenesis. Targeting this protein will be potent choice for antiviral drug designing. The information is much needed to prevent and control developmental abnormality due to ZIKV by modulating these sfRNA interacting host proteins.

## Materials and Methods

### Phylogenetic analysis and sequence alignment

The 3’ UTR sequence of ZIKV retrieved from NCBI database was utilized to construct the phylogenetic tree. The best fit model was identified as K2: Kimura 2-parameter in maximum likelihood method was determined using MEGA7. The evolutionary history was inferred by using the Maximum Likelihood method based on the Kimura 2-parameter model. The tree with the highest log likelihood (-810.97) is shown. The percentage of trees in which the associated taxa clustered together is shown next to the branches. Initial tree(s) for the heuristic search were obtained automatically by applying Neighbor-Join and BioNJ algorithms to a matrix of pairwise distances estimated using the Maximum Composite Likelihood (MCL) approach, and then selecting the topology with superior log likelihood value. The tree is drawn to scale, with branch lengths measured in the number of substitutions per site. The analysis involved 62 nucleotide sequences. All positions containing gaps and missing data were eliminated. There were a total of 414 positions in the final dataset. Evolutionary analyses were conducted in MEGA7.

### ZIKV sfRNA secondary structure prediction

The secondary structures of ZIKV sfRNA were predicted using RNAfold web server (http://rna.tbi.univie.ac.at/cgi-bin/RNAWebSuite/RNAfold.cgi)

### Prediction of host proteins interacting with ZIKV sfRNA

In addition, the sfRNA interacting host protein was predicted using software (http://s.tartaglialab.com/page/catrapid_omics_group) and top 120 interacting proteins were used in the study. Functions of proteins were retrieved from UniProt online source (https://www.uniprot.org)

## Acknowledgements

Author acknowledges Nagalingam R. Sundaresan for providing facility in Department of Microbiology and Cell Biology at the Indian Institute of Science, Bangalore. P.A.D is the recipient of DST-Inspire faculty fellow. P.A.D is supported by research funding from Department of Science and Technology, Government of India DST/INSPIRE/04/2016/001067.

---

**Supplementary Figure 1:**
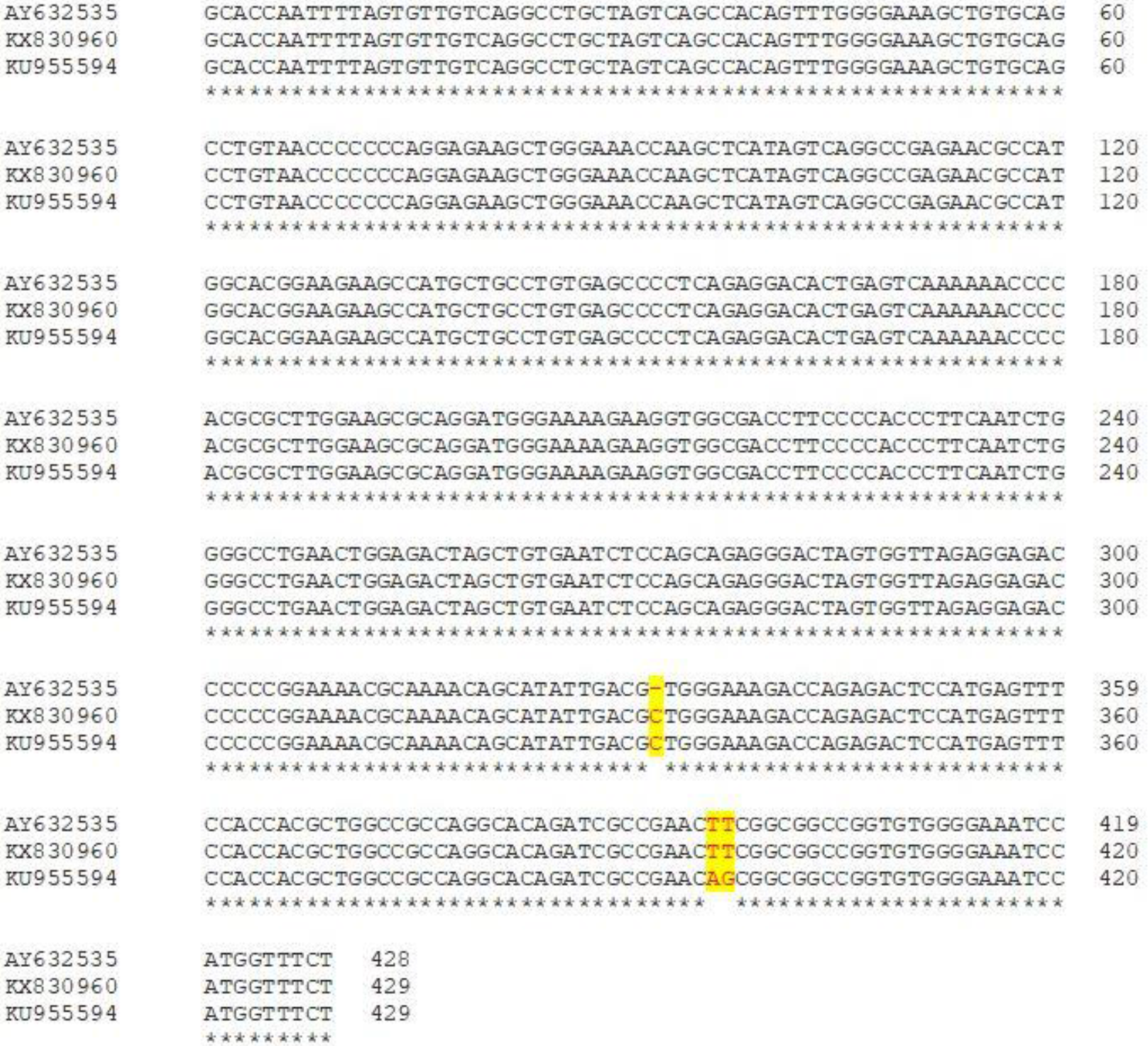
Alignment of MR766 sfRNA. MR766 reference sequence (AY632535), MR766 sequence which has extra nucleotide ‘C’ at the position of 332 (KX830960), clade II MR766 sequences which have extra nucleotide ‘C’ at the position of 332 and mutations of T396A and T397G (KU955594).

**Supplementary Figure 2:**
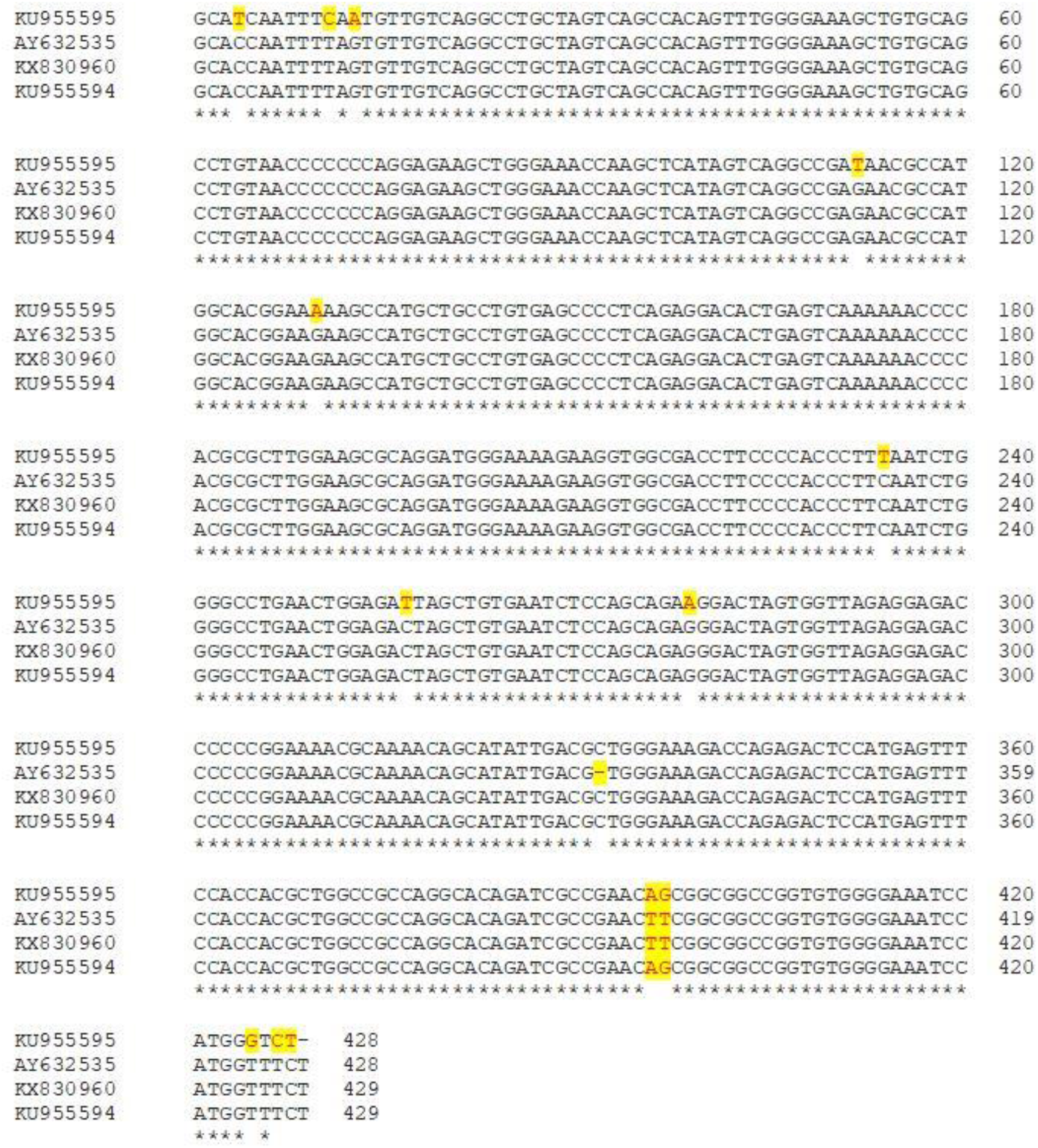
Alignment of African ZIKV sfRNA. MR766 reference sequence (AY632535), MR766 sequence which has extra nucleotide ‘C’ at the position of 332 (KX830960), clade II MR766 sequences which have extra nucleotide ‘C’ at the position of 332 and mutations of T396A and T397G (KU955594) and DAK strain has mutation C4T, T11C, G13A, G112T, G130A, C234T, C257T, G279A, T425G, T427C and C428T.

